# Linking the brain with behaviour: the neural dynamics of success and failure in goal-directed behaviour

**DOI:** 10.1101/2021.05.25.445701

**Authors:** Amanda K. Robinson, Anina N. Rich, Alexandra Woolgar

## Abstract

The human brain is extremely flexible and capable of rapidly selecting relevant information in accordance with task goals. Regions of frontoparietal cortex flexibly represent relevant task information such as task rules and stimulus features when participants perform tasks successfully, but less is known about how information processing breaks down when participants make mistakes. This is important for understanding whether and when information coding recorded with neuroimaging is directly meaningful for behaviour. Here, we used magnetoencephalography (MEG) to assess the temporal dynamics of information processing, and linked neural responses with goal-directed behaviour by analysing how they changed on behavioural error. Participants performed a difficult stimulus-response task using two stimulus-response mapping rules. We used time-resolved multivariate pattern analysis to characterise the progression of information coding from perceptual information about the stimulus, cue and rule coding, and finally, motor response. Response-aligned analyses revealed a ramping up of perceptual information prior to a correct response, suggestive of internal evidence accumulation. Strikingly, when participants made a stimulus-related error, and not when they made other types of errors, patterns of activity initially reflected the stimulus presented, but later reversed, and accumulated towards a representation of the *incorrect* stimulus. This suggests that the patterns recorded at later timepoints reflect an internally generated stimulus representation that was used to make the (incorrect) decision.

These results illustrate the orderly and overlapping temporal dynamics of information coding in perceptual decision-making and show a clear link between neural patterns in the late stages of processing and behaviour.

## Introduction

A primary function of the human brain is to flexibly respond to relevant perceptual information in accordance with current context and task goals. The sound of a phone ringing, for example, should prompt a different response if it is your phone than if it belongs to someone else. This set of complex processes, termed cognitive control, involves interpreting incoming information given the current context to determine an appropriate action (Posner & Presti, 1987; Posner & Snyder, 1975). Cognitive control involves dynamic information exchange at different levels of processing, from perceptual information processing to decision-making and response selection.

Understanding how these different processes unfold could provide a great deal of insight into how the brain achieves goal-directed behaviour.

A large body of neuroimaging research implicates frontoparietal brain regions in goal-directed behaviour, which form a distributed network responsible for cognitive control (Duncan, 2010). This “multiple demand” (MD) network (Duncan, 2010), elsewhere referred to as the cognitive control network (Cole & Schneider 2007), frontoparietal control system (Vincent et al. 2008), or task-positive network (Fox et al. 2005), appears to flexibly represent different types of information depending on task context. For example, activity in these regions encodes task rules (e.g., Crittenden et al., 2016; Woolgar et al., 2015, 2011), and auditory, visual, and tactile stimulus features (Bracci et al., 2017; Jackson et al., 2017; Long and Kuhl, 2018; Woolgar and Zopf, 2017; for a review see Woolgar et al., 2016). These regions particularly encode task elements that are demanding (Woolgar, Afshar, et al., 2015; Woolgar et al., 2011) or at the focus of attention (Jackson et al., 2017; Jackson & Woolgar, 2018; Woolgar, Williams, et al., 2015). Activity in some of these regions has also been causally implicated in selectively facilitating coding of task-relevant information (Jackson et al., 2021). This lends support to the possibility that flexible responses within the MD regions play a causal role in goal-directed behaviour (e.g., Duncan et al., 2020; Woolgar et al., 2019, 2018, 2010).

A characteristic feature of cognitive control is that it dynamically changes in response to task-relevant information. Research using fMRI has yielded insight into the brain networks involved in goal-directed behaviour, but the slow nature of the blood-oxygen-level response has limited the exploration of the corresponding dynamics. Time-resolved neuroimaging methods such as magnetoencephalography and electroencephalography (M/EEG) have been more fruitful in understanding how cognitive control unfolds over time (for review, see Gratton et al., 2017). For example, conflict-related processing involving incongruent task features elicits a larger evoked response than a congruent condition approximately 200-400ms after stimulus onset, which has been linked to activity within the anterior cingulate (Folstein & Van Petten, 2008), and task-switching involves a larger parietal positivity around 300ms relative to task repeats (Karayanidis et al., 2010). The newer method of multivariate pattern analysis (MVPA) in conjunction with M/EEG allows further insight into processing dynamics underlying cognitive control. MVPA uses pattern classification approaches applied to neuroimaging data to show what information is being coded within the brain (Haxby, 2012; Haxby et al., 2001). Time-resolved MVPA has been used to characterise how information coding changes over time (Carlson et al., 2011; Hebart & Baker, 2018). For example, Hebart and colleagues (2018) had participants perform different tasks on visual object stimuli while measuring MEG, and showed that task-relevant object features were enhanced at late stages of processing, more than 500ms after the stimulus was presented. Other work has shown clear progression of task relevant information during complex tasks, with different dynamics for features such as stimulus, task and response (Hubbard et al., 2019; Kikumoto & Mayr, 2020; Wen et al., 2019). This line of research has also highlighted the importance of combining relevant task information for successful behaviour (Kikumoto & Mayr, 2020). These MVPA studies provided great insight into the neural dynamics of goal-directed behaviour, but all used designs where the task cue was presented prior to the target, allowing participants to prepare for the task in advance. Additionally, these studies focused on stimulus-aligned neural responses. It seems likely that tracking the dynamic coding of relevant task features relative to both stimulus onset and response, using a task that induces more flexible behaviour, might elucidate stronger links between dynamic neural responses and goal-directed behaviour.

Decades of neuroimaging research have focused on the neural correlates of behaviour, but assessing whether particular patterns of brain activity are *necessary* for behaviour has presented a challenge. In MVPA, a classifier algorithm is trained to distinguish between conditions using patterns of neural data from multiple trials of each condition. If a classifier can predict the conditions of new neural data better than chance, this demonstrates that the patterns of activity in the data must contain, or represent, information about the different conditions. However, the conclusion that decodable patterns represent “information” has been questioned on theoretical grounds (de-Wit et al., 2016): just because information is decodable using machine learning does not necessarily mean it is used by the brain to generate behaviour. This awareness has lead researchers to push for more explicit links between MVPA patterns and behaviour, for example, comparing details of patterns to reaction times or accumulation rates in models of behaviour (Grootswagers et al., 2018; Ritchie et al., 2015; Ritchie & Carlson, 2016).

Exploring how information coding changes when participants make errors is another way to establish how behaviourally meaningful patterns of activation are. For example, Williams et al (2007) demonstrated that multivariate fMRI patterns in lateral occipital cortex, but not those in early visual regions, reduced to chance when participants made errors on a shape discrimination task, indicating that patterns in early visual cortex were not directly read out in behaviour. In another study, participants performed a scene classification task (Walther et al., 2009), and classifier prediction error patterns correlated with the types of errors in behaviour within high level object and scene-specific brain regions, but not within early visual cortex. Using magnetoencephalography (MEG), we have recently shown that this logic can even be used to predict behavioural errors before they occur (Karimi-Rouzbahani et al., 2021).

A stronger requirement for a behaviourally meaningful pattern of activity is that it should not just change on error, but change to something that predicts the particular error to be made (Woolgar et al., 2019). We tested this in fMRI, and found that patterns of activation in frontoparietal cortex indeed reversed on error, such that patterns of activation on error trials represented information that was not presented to the participant, in a manner that was diagnostic of the particular behavioural error they made (Woolgar et al., 2019). In that study, participants performed a difficult response-mapping task. When participants made a rule error, MD patterns of activity reflected the *incorrect* rule, and when participants made other errors, MD patterns of activity reflected the *incorrect* stimulus (Woolgar et al., 2019). Within visual cortex, in contrast, there was no evidence of relevant information (correct or incorrect) during errors. Thus, some multivariate patterns appear to be more directly relatable to behaviour that others, and there is a tight link between frontoparietal activity patterns and behavioural outcome.

In the current study, we used MEG and MVPA to examine the dynamics of this effect, asking whether information coding through the course of a trial was equally associated with behaviour. We aimed to (1) characterise the neural dynamics of multiple types of task relevant information, and (2) examine their relationship to behaviour over time. Participants performed a difficult response-mapping task which required different responses to a target stimulus depending on the current rule. To determine what aspects of this representation could be directly linked to behaviour, we examined information coding on *incorrect* trials: when the wrong rule was applied or when there were errors in perception. We found a clear progression in onset of information coding, such that stimulus features are evident shortly after stimulus onset, followed by abstract rule coding and then the response, with the information about each task feature accumulating up to the time of response. When participants made stimulus errors, stimulus information was initially coded veridically but later accumulated in the opposite direction, towards a representation of the incorrect stimulus. By contrast, stimulus information was encoded correctly when participants made rule errors. The data reveal the dynamics with which information coding in the brain can be tightly linked to participant behaviour.

## Methods

All data, stimuli and code are available on the Open Science Project at https://osf.io/2nwhr/.

### Participants

Participants were 22 healthy adults (14 females, 8 males, age range 18-38 years) with normal or corrected-to-normal vision recruited from Macquarie University. This study was approved by the Macquarie University ethics committee and informed consent was obtained from all participants. Participants took part in two sessions: a 1-hour behavioural session, and a 2-hour MEG session, on separate days. They were compensated $15 for the behavioural session and $40 for the MEG session. For two participants, initial photodiode inaccuracies meant that the timing for 2 and 5 trials respectively was not adequately marked, so these trials were excluded from analyses. Data from an additional two participants (2 males) were collected and excluded: both participants had very few stimulus position errors during the MEG session (<10), and for one of the participants there was a recording error such that MEG data was recorded for only 680 out of 800 trials.

### Design and procedure

Participants learned to apply two difficult response-mapping rules regarding the position of a target stimulus. The target was a grey square approximately 2x2 degrees of visual angle that appeared in one of four positions. All positions were equidistant from fixation at an eccentricity of 4 degrees of visual angle. Within the left and right visual fields, the two possible target locations overlapped by 60% horizontally and 65% vertically to create a high degree of position uncertainty (Figure 1a). Participants had to respond to the position of the stimulus using two possible response-mapping rules (Figure 1a). The two rules each comprised 4 unique position transformations and were mirror images of each other. The colour of a central fixation square acted as a cue for the rule. There were two cues per rule, in order to dissociate neural responses to cues from the neural responses to rules (e.g., blue and yellow = Rule 1, pink and green = Rule 2; counterbalanced across participants). Participants responded by pressing one of four response keys with their right hand. The stimuli were presented using PsychToolbox in MATLAB.

**Figure 1.**
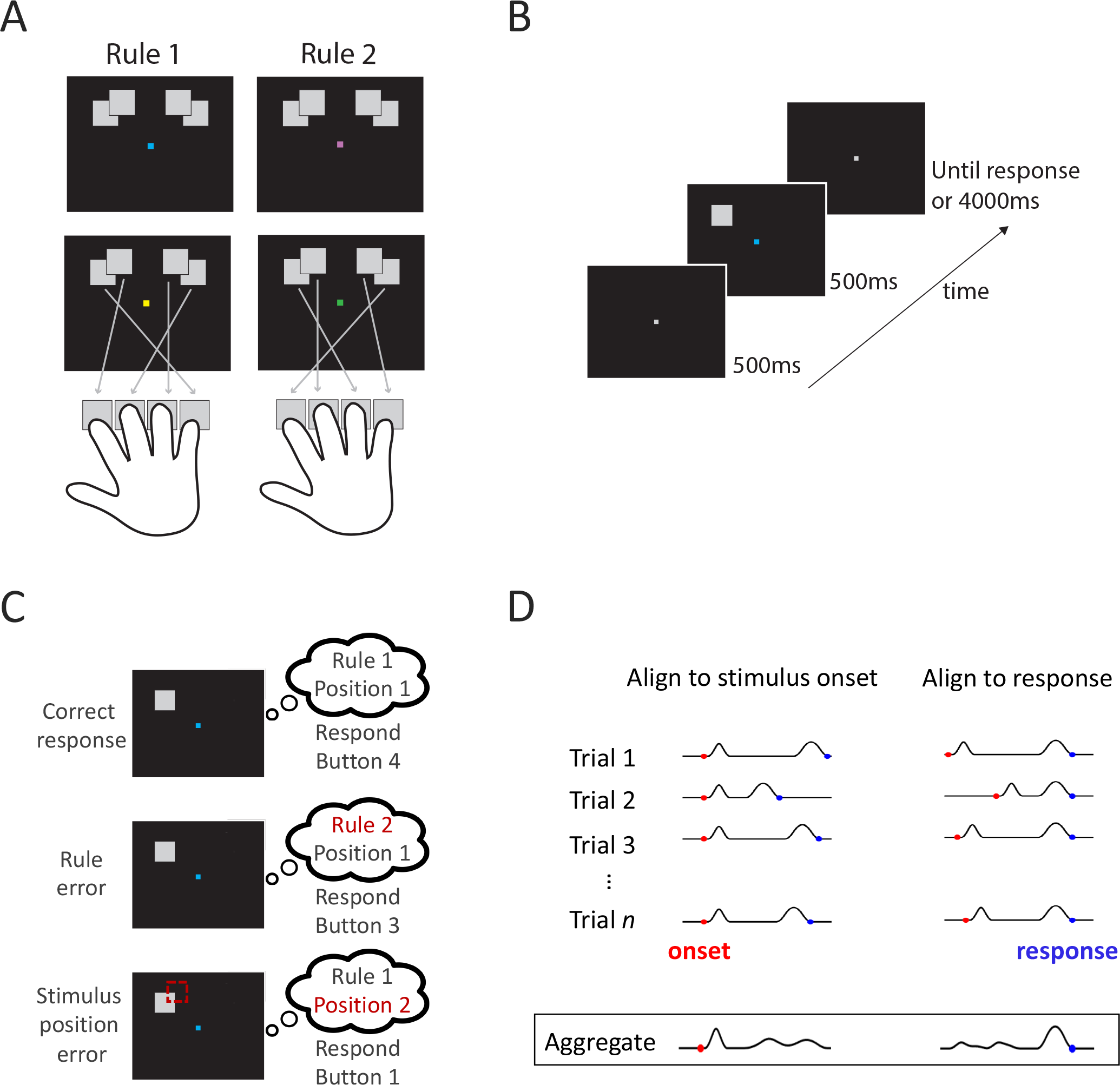
Experimental design. A) Response mapping rules. Participants had to indicate the position of a target stimulus which appeared in one of four possible locations. There were two cues per rule, designated by blue/yellow and green/pink squares at fixation. The button press associated with each position is indicated by the specific rule. B) Trial timeline. After a fixation screen, the target stimulus and coloured fixation cue appeared simultaneously, and participants had to apply the correct response-mapping rule using a button press. C) Behavioural response types. In this example, the stimulus was in Position 1 and the rule was Rule 1 (blue cue), so the correct response was Button 4. A rule error occurred if the rule was mistaken for Rule 2, leading to a response of button 3. A stimulus position error occurred if the position was mistaken to be Position 2, leading to a response of button 1. D) Depiction of MEG data collation: data aligned to stimulus onset (left) or response time (right). The temporal dynamics of stimulus-related and decision-related neural responses vary across trials, with different processes aligned with onset and response. Aligning the MEG data to stimulus onset versus response highlights different neural stages on the aggregate of all trials, even though the content of each trial is identical.

The stimulus-response rule mappings were designed to distinguish correct responses and specific types of errors (Figure 1c). An error was considered a *rule error* when the button press response reflected the combination of the correct stimulus position with the wrong rule. In contrast, a *stimulus error* was defined as a button press response consistent with the combination of the adjacent perceptually confusable position with the correct rule. For example, under Rule 1, if the stimulus appeared on the far left, the correct response would be button 4, a rule error (i.e., using Rule 2 applied to the correct position) would lead to a button 3 response, a stimulus error (i.e., using Rule 1 correctly but confusing the stimulus with the other left position) would lead to a button 1 response, and confusing both the rule and the stimulus led to a button 2 response.

### Training session

Participants learned each rule in a separate session outside the MEG. They were trained to perform the task using increasingly difficult blocks of trials (see below). Feedback was given on every trial. For every incorrect response, participants were shown the correct response.

Initially, stimuli were presented in non-overlapping positions (i.e., further apart than the final paradigm) so there was no position uncertainty. Stimuli were presented on the screen until a response was made (i.e., not time-limited). In the first block, participants learnt the first rule (Rule 1 or Rule 2, counterbalanced across participants). Each stimulus position was shown with its associated response four times (16 trials), and participants had to press the appropriate button for each stimulus. In each trial, cue colour was chosen randomly from the two possible cues for that rule. The second block followed the same protocol, but for the other rule (Rule 1/Rule 2). In the third block, participants had to perform the task by implementing both rules, but still with well separated stimuli. In the fourth block, the stimuli were presented in their final, overlapping experimental positions. Finally, in the fifth block, the stimuli were presented with the same procedure as the final experimental paradigm: the stimuli were overlapping and were presented for only 500ms. Blocks 3, 4 and 5 contained 32 trials each, consisting of 2 repeats of each cue and stimulus position, randomly ordered. In all blocks, participants had to perform at 60% accuracy or above to progress to the next block type. Blocks were repeated if they did not reach this threshold. On average, participants completed 8.61 training blocks (*SD* = 2.46). Block 3 was most often repeated (*M* = 3.09 repeats).

### Experimental session

In the second session, participants performed the task while their neural activity was recorded using MEG. At the start of each block, participants were shown a graphical depiction of the rules for at least 2 seconds. When they were ready, they pressed a button to begin the block. In each trial, participants were shown a grey fixation marker for 500ms, and then the square target stimulus and coloured rule cue were presented for 500ms (Figure 1b). The participants were instructed to respond as quickly as possible without sacrificing accuracy. After they responded, there was an inter-trial interval of 1000ms before the next trial started. There were 10 blocks of 80 trials, each containing 5 trials per stimulus and cue colour combination. Within each block, the order of the trials was randomised. Instead of feedback on every trial, like in training, participants were given feedback about their mean accuracy and reaction times at the end of each block.

### MEG acquisition

MEG data were collected at Macquarie University in the KIT-MQ MEG facility with a whole-head supine Yokogawa system containing 160 gradiometers (Kado et al., 1999). The participant’s head was fitted with a cap containing five marker coils. The head shape and position of the marker coils was marked using a Polhemus digitisation system. Once inside the MEG, the position of the marker coils was measured to ensure the MEG sensors had good coverage over the participant’s head. Marker position measurements were repeated halfway through the experiment and at the end of the session. Raw MEG data were collected at 1000 Hz with online 0.03 Hz highpass and 200 Hz lowpass filters.

Stimuli were projected onto the ceiling of the magnetically shielded room. Stimulus timing was measured using a photodiode placed on the projection mirror and marked in an additional channel in the MEG recording. Participants indicated their response using a 4-Button Fiber Optic Response Pad (Current Designs, Philadelphia, USA). Response timing was marked in the MEG recording using a parallel port trigger.

### MEG data analysis

MEG data were analysed using multivariate decoding, which is very sensitive to reliable effects in the data and resistant to artefacts such as eye blinks that are not consistent across time and condition (Carlson et al., 2020; Grootswagers et al., 2017). Due to the robustness of decoding to such artefacts, data were minimally preprocessed using EEGLAB (Delorme & Makeig, 2004). Data were filtered using a Hamming window FIR filter (default EEGLAB filter pop_eegfiltnew) with highpass of 0.1Hz and lowpass of 100Hz, and then downsampled to 200 Hz before epoching. For separate analyses, trials were epoched relative to stimulus onset, marked by the photodiode, and to the button press response, marked by the parallel port trigger.

Data were analysed using time-resolved classification methods (e.g., Carlson et al., 2020; Grootswagers et al., 2017) and implemented using the CoSMoMVPA toolbox (Oosterhof et al., 2016). For each time point, data were pooled across all 160 MEG sensors, and we tested the ability of a linear discriminant analysis (LDA) classifier to discriminate between the patterns of neural responses associated with the different conditions. Trials were divided according to their associated behavioural responses: correct trials, rule errors and stimulus position errors (Figure 1C). The classifiers were always trained on correct trials. To ensure that there were equal numbers of trials for each condition, correct trials were subsampled to be equal for each position and rule combination for each block per participant. To ensure adequate trial numbers for each of the analyses, blocks with fewer than two trials per rule x position combination were excluded; this amounted to 9 excluded blocks in total across 8 participants, with the remaining 14 participants having all blocks included. The total number of selected trials per participant was *M* = 437.09 (min = 280, max = 600).

### Temporal dynamics of stimulus, cue, rule and response coding

We performed pattern classification analyses to determine the time points at which stimulus position, cue, rule and response representations emerge in the brain. First, we decoded stimulus position by comparing neural representations of the inner two stimulus positions (Positions 2 and 3) to those of the outer two stimulus positions (Positions 1 and 4). Separating position in this manner meant that motor responses and rule were balanced across the two position conditions and could not drive the classification results, ensuring we are detecting information related to stimulus position.

Next, we assessed the time course of rule coding by training a classifier to distinguish between Rule 1 and Rule 2. In having two colour cues per rule, this analysis focused on rule coding over and above the physical properties of the cues (Rule 1 (blue and yellow cues) versus Rule 2 (pink and green cues).

We can also, however, decode cue coding separately from rule coding. To assess how cue decoding differed from rule decoding, we decoded between the two cues per rule (i.e., blue versus yellow colour cue for Rule 1, and pink versus green for Rule 2). Cue coding was quantified as the mean of the two pairwise analyses.

As a final analysis, we decoded motor response by comparing the inner two button presses to the outer two button presses. This comparison ensured that stimulus position and rule were balanced within each class, so that the classifier would be driven by the motor response alone.

For each decoding analysis, classification analyses were performed using a leave-one-block out cross-validation approach. This resulted in 10-fold cross-validation for participants with no excluded blocks (*N* = 14). The remaining participants used 9-fold (N = 7) and 8-fold cross-validation (*N* = 1). For all decoding analyses, chance performance was 50%.

### Error representations

The next set of analyses focused on decoding neural activity when participants made errors, to explore the relationship between patterns of activity and behaviour. To investigate the representation of rule and stimulus errors, we trained the classifier on the correct trials and tested on the error trials. This allowed us to decode what information was present in the patterns of response across sensors when participants made different kinds of mistakes. Specifically, the analyses assessed whether the error patterns resembled the *correct* stimulus and rule patterns (above chance decoding), or the neural patterns associated with the *incorrect* stimulus and rule (below chance decoding). Note that in this approach, **below chance classification is meaningful**: it indicates the representation of the pattern that is instantiated when the other (incorrect in this case) rule or stimulus position is encoded.

We performed error decoding for stimulus position and rule information. In a comparable procedure to the correct trial analysis, we used leave-one-block out cross-decoding analyses. In each fold, the classifier was trained on *correct* trials from all but one block, and tested on all *error* trials across the whole session. This ensured the same training data (and thus decoding models) were used as in the correct trial analyses, but allowed well-characterised results for the relatively small number of error trials. Participants made an average of 5.71% rule errors and 9.82% stimulus position errors (Figure 2A). Table 1 shows the mean number of trials used for stimulus and rule decoding per trial response type (correct, stimulus error, rule error).

**Figure 2.**
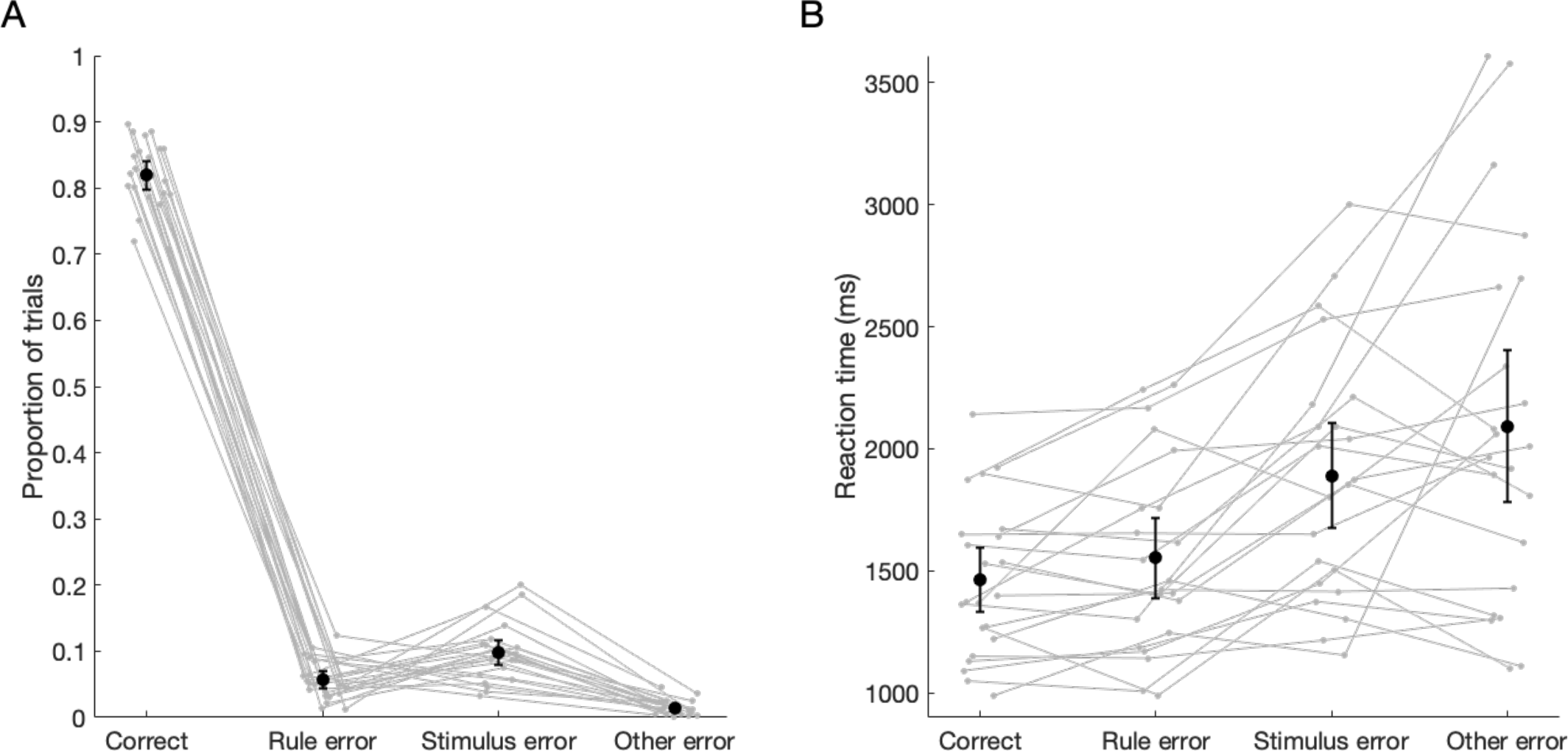
Behavioural results in the MEG session (*N* = 22). A) Proportion of trials, and B) Median reaction time for correct, rule errors, stimulus errors and other errors. Grey lines denote individual participants and black markers denote group means. Error bars are 95% confidence intervals across participants.

**Table 1.** Mean number of trials used for stimulus and rule decoding analyses per participant. Group means (standard deviation) for trial numbers are presented according to the two different stimulus classes (inner or outer positions; regardless of rule) and two rules (regardless of stimulus position). Trials are split by behaviour: correct response, stimulus error and rule error. Decoding models were always trained using correct trials.

|  | Correct | Stimulus error | Rule error |
| --- | --- | --- | --- |
| Inner positions | 218.55 (46.00) | 30.00 (29.35) | 17.55 (9.14) |
| Outer positions | 218.55 (46.00) | 48.50 (26.05) | 28.14 (17.94) |
| Rule 1 | 218.55 (46.00) | 38.14 (16.09) | 22.18 (13.29) |
| Rule 2 | 218.55 (46.00) | 40.36 (21.44) | 23.50 (12.19) |

### Exploratory Searchlight analysis

To illustrate the spatial extent of the MEG signal containing relevant task-related information, we applied searchlight decoding. For each sensor in turn, we defined a searchlight consisting of that sensor and its immediate neighbours (mean 5.6 sensors per searchlight). We then ran the same decoding schemes as on the whole head (above) in each of these searchlights. Decoding accuracy for each searchlight was plotted on the central sensor, resulting in a head map of decoding accuracies, showing the regions containing task-related information at a given time. For each task feature and condition (e.g., rule 1 vs rule 2 on stimulus-aligned correct trials), searchlight results were plotted using 20 ms time windows of interest centred around representative timepoints: 150ms and 1000ms for the onset-aligned analyses, and -600ms and -200ms for the response-aligned analyses.

### Statistical testing

To assess performance of the classifier, we used Bayesian statistics to determine the evidence that decoding performance was different from chance (Dienes, 2011, 2016; Jeffreys, 1961; Rouder et al., 2009; Teichmann et al., 2021; Wagenmakers, 2007). A Bayes factor (BF) is the probability of the data under the alternative hypothesis relative to the null hypothesis. In all analyses, the alternative hypotheses of above- and below-chance (50%) decoding were tested using the ‘BayesFactor’ package in R (Morey et al., 2018). Bayes Factors were calculated using a JZS prior, centred around chance decoding of 50% (Rouder et al., 2009) with default scale factor of 0.707, meaning that for the alternative hypotheses of above- and below- chance decoding, we expected to see 50% of parameter values falling within -.707 and .707 standard deviations from chance (Jeffreys, 1961; Rouder et al., 2009; Wetzels & Wagenmakers, 2012; Zellner & Siow, 1980). A null interval was specified as a range of effect sizes between -0.5 to 0.5.

In accordance with the Bayes Factor literature, we did not make a correction for multiple comparisons for the large number of time points tested (Dienes, 2016, 2011; Świątkowski and Carrier, 2020; Teichmann et al., 2021). Bayes Factors assess the strength of evidence for the null hypothesis versus the alternative hypothesis; here, they are used to directly test the evidence for above-chance (or below-chance) decoding and the evidence that decoding is equivalent to chance. Thus, we assess the magnitude of evidence in either direction, rather than a probability of the observed data occurring by chance as in traditional null hypothesis testing. BF > 3 is typically considered evidence for the alternative hypothesis, and BF < 1/3 as evidence in favour of the null hypothesis (Jeffreys, 1961; Wetzels et al., 2011). When the magnitude of Bayes Factors are interpreted at face value, rather than applying a threshold for “significance”, there is no need to correct for the number of tests (time points) (Teichmann et al., 2021). Additional tests provide additional evidence, and can be interpreted as such without correcting for multiple comparisons (Dienes, 2011, 2016). Accordingly, we do not interpret high Bayes Factors at isolated time points as evidence for decoding, rather we assess the pattern of evidence through time in support of the null or alternative hypotheses. Single time points are not considered to provide substantial evidence if neighbouring time points support the opposite hypothesis. We will refer to periods of time with sustained evidence for the alternative hypothesis as times when information “could be decoded”, indicating information was represented in the brain.

## Results

### Behavioural results

All participants performed above 60% on the final block of the response-mapping task in the training session and therefore participated in the experimental MEG session. In the MEG session, participants performed well above chance (*M* = 81.92%) but still made both rule errors (*M* = 5.71%) and stimulus position errors (*M* = 9.82%; Figure 2a). There were very few other errors (*M* = 1.43%) or trials with no response (*M* = 1.13%). Reaction times were slower for stimulus error trials than rule error and correct trials (Figure 2b).

### Temporal dynamics of goal-directed behaviour

First, we investigated neural coding during correct trials by decoding different task-related information from the MEG signal when each trial was aligned to stimulus onset (analogous to classic event-related analyses). We then realigned the MEG signal of each trial to the response and performed the same decoding analyses (see Figure 1D for depiction of realignment). This gives us a unique insight into the time course of the processing stages during goal-directed behaviour.

Time-resolved decoding performed relative to stimulus onset revealed a progression of relevant information over time (Figure 3a). Stimulus position information was represented in the neural signal from approximately 75ms after the stimulus appeared, with a double-peak response. Cue information could be decoded from 170ms, and the dynamics were similar for the blue versus yellow (53.34% at 170ms) and green versus pink (51.86% at 170ms) cue colour decoding (mean presented in Figure 3a), indicating the decoding reflects general cue information rather than being specific to one set of cues. The timing of stimulus and cue information was thus consistent with early visual stages of retinotopic position (Battistoni et al., 2020; Carlson et al., 2011) and colour processing (Teichmann et al., 2019), respectively.

**Figure 3.**
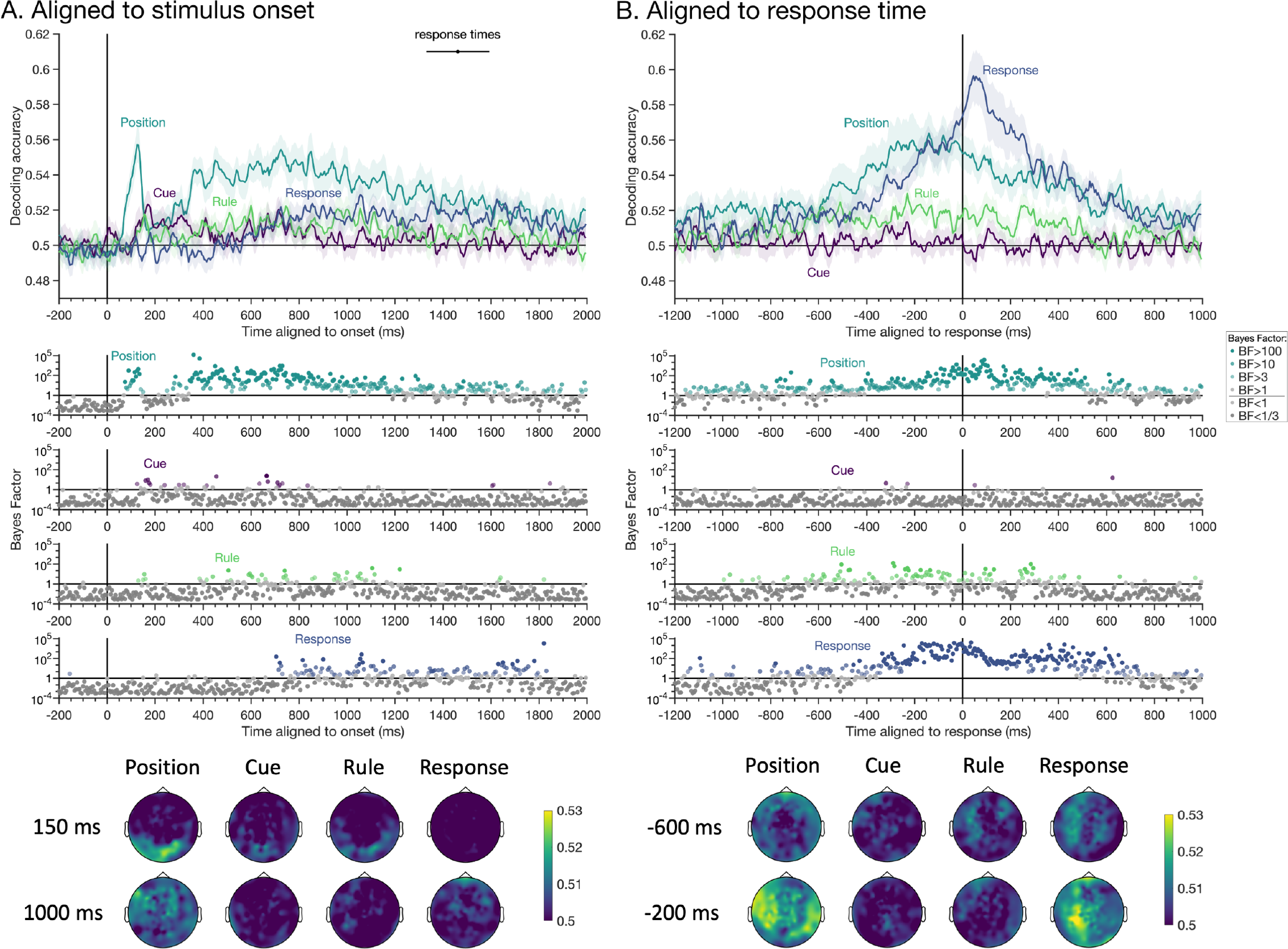
The temporal dynamics of correct stimulus position, cue colour, rule and response information coding. A) Decoding analyses conducted relative to stimulus onset. B) Decoding analyses conducted relative to response time. Shaded areas show standard error across participants (*N* = 22). Decoding accuracy is smoothed with a 20ms window for visualisation. Line at top of plot A marks the mean and 95% confidence interval of median response times per participant. Bayes factors (BF) for above-chance decoding are displayed below the x-axes for every time point using a log scale and colour coded according to the evidence for above chance decoding (see inset). Headmaps depict sensor searchlight decoding results for each task-related feature at representative time periods. The colour bar indicates decoding accuracy per sensor searchlight, calculated as the mean of a 20 second time window centred on the time of interest.

Rule information was briefly evident at about the same time as the cues (around 150ms), but also prominent from approximately 400 to 1200ms, likely coinciding with higher-level cognitive stages of processing. Rule information was quite low in general. One possibility is that there was a carryover effect such that the rule type on a given trial subsequently affected rule coding on the next trial. In an additional exploratory analysis, we assessed rule decoding separately for rule switch and rule repeat trials by training rule 1 versus rule 2 on correct trials (as in the original analysis) and testing on repeat and switch trials separately. We found that rule decoding exhibited similar dynamics regardless of the rule on the previous trial, with slightly higher decoding on switch trials (see OSF repository). Thus, inertia from the previous trial did not appear to cause a switch cost in the representation of rule information in the brain.

Finally, we assessed the temporal dynamics of response information. The button press response was represented from about 715ms, corresponding with reaction times on some of the faster trials. Together, decoding accuracy peaked at 125ms for stimulus position, 175ms for cue colour, 600ms for rule and 1055ms for response button, showing a clear progression in information processing through time.

Searchlight decoding results showed that position, cue and rule information at 150ms localised to posterior regions of the brain (Figure 3A, bottom). At a later time period, 1000ms, position and response information were more frontal and diffuse across the MEG sensors. Rule information was much lower in general but again showed a dissociation for the early versus late time points, with frontal and central topography at the later time period.

As is typical in difficult tasks, there was a wide variation in response times across trials and participants, indicating that the dynamics of high-level task-related processes such as decision-making vary trial to trial with respect to stimulus onset. Time-resolved decoding relies on processes occurring at the same time across trials, so this temporal jitter can mask results (Vidaurre et al., 2018). In order to capture processes that are more closely aligned with the response, we next realigned the MEG data to the response time (Figure 1D) and performed the same decoding analyses. The temporal dynamics of relevant task-related information were markedly different compared with onset-aligned results (Figure 3B). Notably, cue information could no longer be reliably detected, presumably because cue colour representations were transient and tightly stimulus-locked since, by design, the cue-distinctions were irrelevant as soon as rule information could be extracted from them. In contrast, decoding of stimulus position coding was evident more than 1000ms before the response, and rule coding was evident more than 600ms before the response, with evidence for stimulus processing earlier than rule processing. Motor response coding was sporadically present from more than 1000ms before the response, but was sustained from around 485ms prior to when the response was made. Response coding peaked after the response was given, potentially reflecting the contribution of somatosensory feedback from the different button presses. Response coding 200ms before the response was associated with highest decoding over central-left sensors, which would be consistent with motor and somatosensory cortex activity associated with a right-hand response. Interestingly, the representation of stimulus position and response information appeared to ramp up before the response, plausibly reflecting the accumulation of evidence leading to a decision.

### Error representations

We were particularly interested in understanding whether and how the task-related information we can decode with MVPA is related to participant performance. Specifically, we investigated how information coding changes when an error is made. Recall that our design explicitly allows us to identify the likely source of the error based on the behavioural response (Fig. 1C). We assessed stimulus and rule information in the neural signal when participants made stimulus errors versus rule errors. Based on our previous work with fMRI (Woolgar et al., 2019), we hypothesised that the brain would represent the incorrect stimulus prior to a stimulus error, and the incorrect rule prior to a rule error. Classifiers were trained to classify stimulus position and rule using correct trials and tested on incorrect trials. Therefore, for each time point on each error trial, the analysis reveals whether activation patterns were more similar to the usual patterns for the presented rule and stimulus (“correct” rule and stimulus) or the alternate one (“incorrect”) corresponding to the participant’s decision (as shown by the behaviour response).

### Stimulus decoding – aligned to stimulus onset

In this analysis, we looked at how stimulus position was coded on error trials. We found that when participants made <u>rule</u> errors, in which behaviour suggested that the stimulus was encoded correctly but the incorrect rule had been used, there was sustained stimulus decoding with similar dynamics to that on correct trials (Figure 4A; blue line). Stimulus coding on <u>stimulus</u> errors, however, was present only transiently at 335 ms after which coding attenuated (Figure 4A; green line). After 495 ms, there was substantial evidence that stimulus coding was higher when participants made errors based on applying the wrong rule than when the response suggested they had misperceived the stimulus. This indicates that when participants made stimulus errors, correct stimulus information was lost.

**Figure 4.**
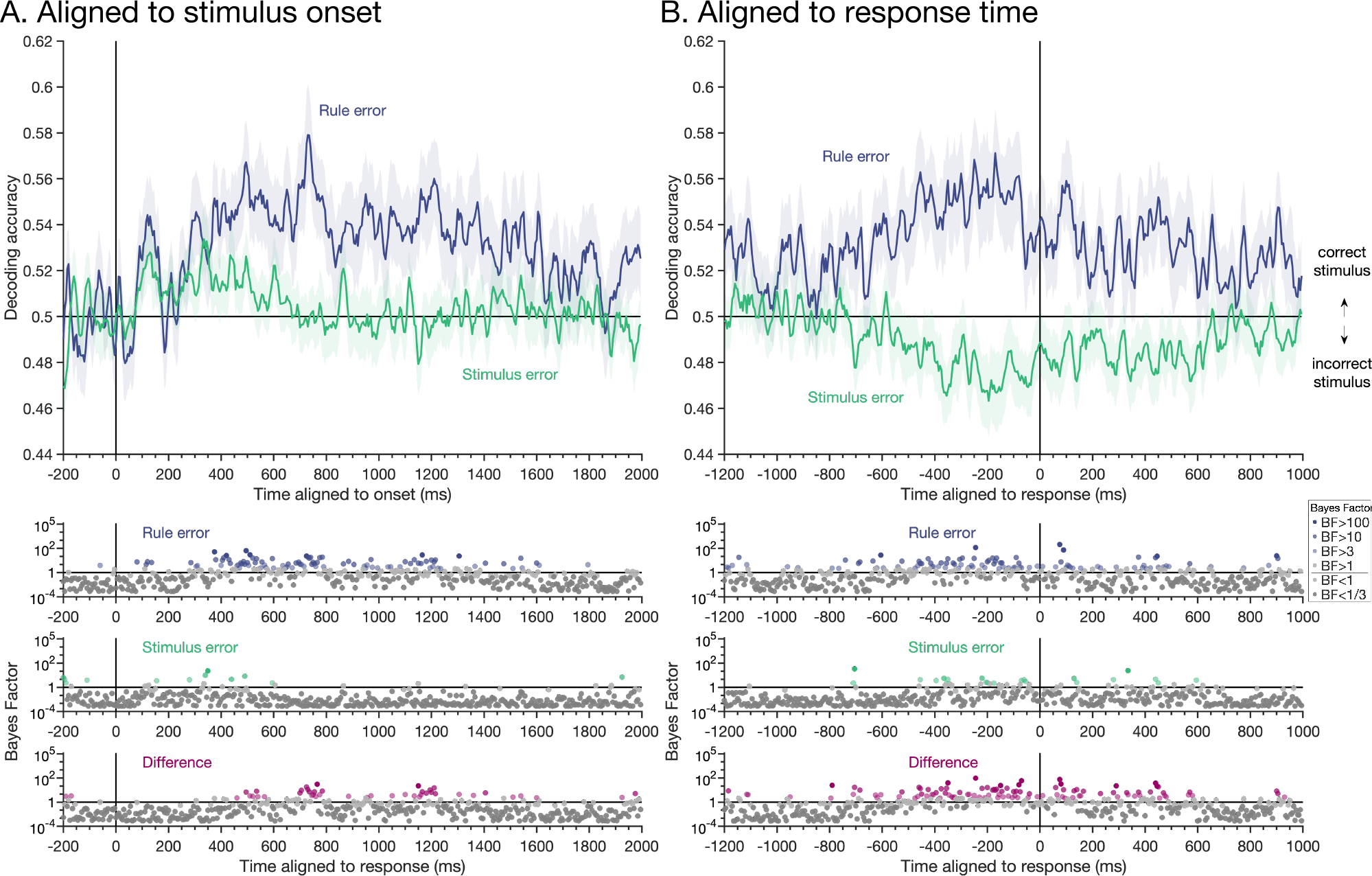
Stimulus position decoding on rule and stimulus error trials. A) Decoding analyses conducted relative to stimulus onset revealed initial stimulus coding on both rule and stimulus error trials, with sustained stimulus coding on rule error trials (similar to correct trials, Figure 3), but no stimulus coding at later timepoints on stimulus error trials. The interaction, shown by BF difference (pink) confirmed that at later timepoints there was more evidence for stimulus coding on rule errors than stimulus errors. B) Decoding analyses conducted relative to response time revealed evidence for correct stimulus coding on rule error trials and evidence for *incorrect* stimulus coding on stimulus error trials, evident as below-chance decoding accuracy. Decoding accuracy is smoothed with a 20ms window for visualisation. Bayes factors are shown on a log scale and colour coded according to amount of evidence.

### Stimulus decoding - aligned to response time

Next, we asked the same question but with the data re-aligned to the response time. On <u>rule</u> errors, there was a gradual ramping up of stimulus coding in the lead up to the response (Figure 4B; blue line), as we had observed for correct trials (Figure 3B). In contrast, on <u>stimulus</u> errors, activation patterns ramped towards the patterns encoding the *incorrect* stimulus, as indexed by below-chance decoding from approximately 355ms before the response (Figure 4B; green line). Given that the correct stimulus had been encoded in the early part of these same trials (Figure 4A), this suggests an evolution of information coding towards the incorrect stimulus decision.

Stimulus decoding accuracy on rule errors was higher than that on stimulus errors for the bulk of the epoch, particularly from about 795ms before the response to 600ms after the response.

Together, this finding shows that stimulus coding in the latter part of the trial reflected the decision ultimately made by participants, rather than the stimulus presented.

### Rule decoding

Next, we asked whether the representation of task *rule* in the correct trials would also generalise to error trials. However, there were only very brief periods of evidence for rule information coding on rule errors and stimulus errors, whether we aligned the MEG data to the stimulus onset (Figure 5A) or response (Figure 5B). There was also no difference in rule coding between error types. For onset-aligned analyses, Bayes Factors indicated evidence for above-chance decoding on rule and stimulus errors for some time points, but it was not sustained. There were also some brief periods of below-chance decoding on rule errors for response-aligned analyses, which would indicate coding of the incorrect rule, consistent with behaviour, but this did not reach our interpretation levels for BFs (2 consecutive timepoints BF>3). Overall, rule information that had been (weakly) present on correct trials was largely absent on both types of behavioural error. Moreover, the *reversal* in coding – in this case coding of the incorrect rule – was not evident as it was for stimulus coding.

**Figure 5.**
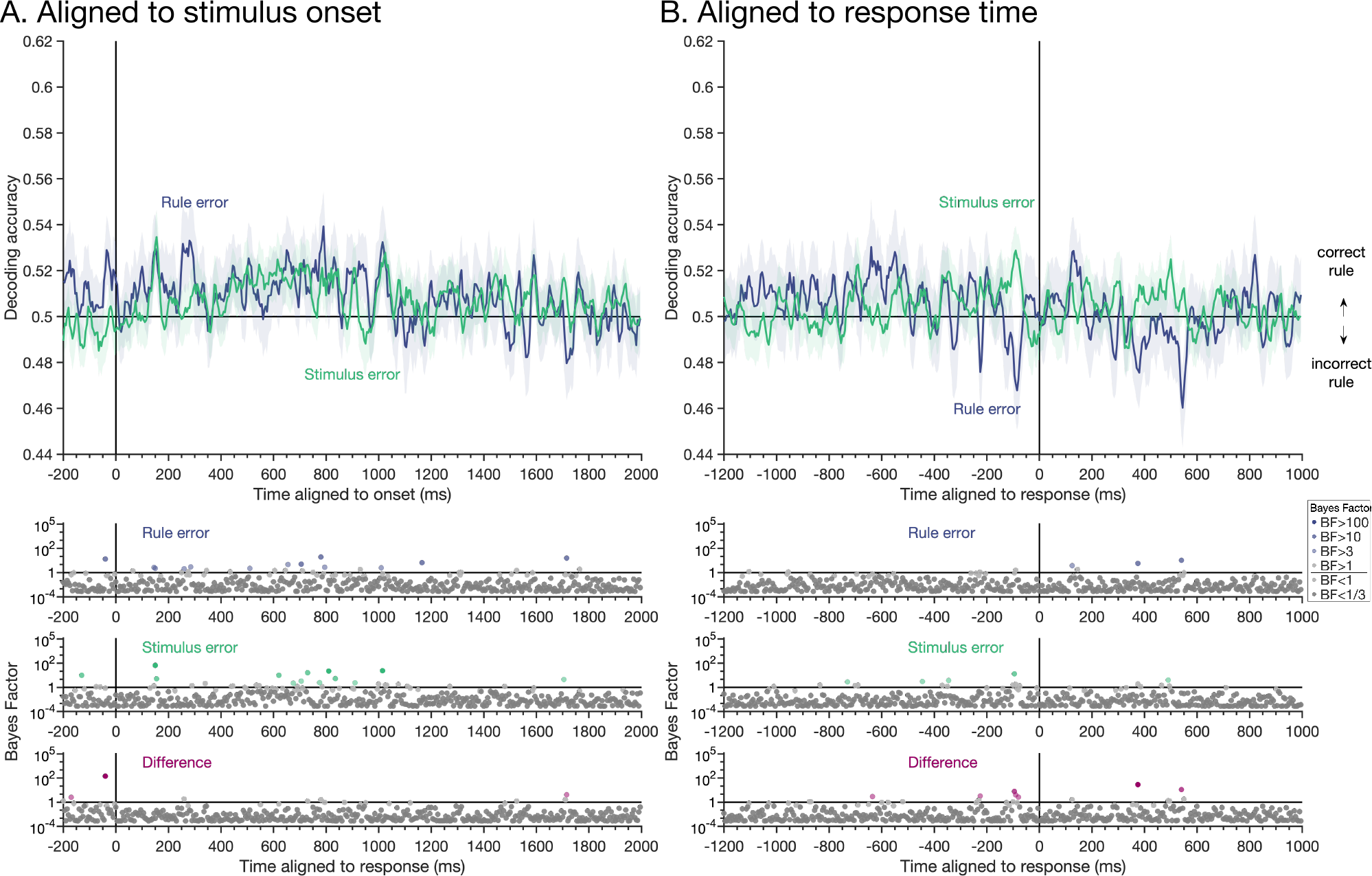
Rule decoding on rule and stimulus error trials. A) Decoding analyses conducted relative to stimulus onset. B) Decoding analyses conducted relative to response time. Decoding accuracy is smoothed with a 20ms window for visualisation. There was substantial evidence for the null (i.e., that rule could not be decoded) on both stimulus error and rule error trials, indicating the rule coding did not reverse for either type of error.

Taken together, the MEG decoding results show that on correct trials, all task-relevant aspects (stimulus position, cue colour, rule, response) could be decoded. The dynamics of coding varied such that analyses revealed stimulus-locked coding of perceptual features (stimulus position, cue) early in the time course, and analyses aligned to the response revealed coding of relevant task aspects (position, rule) and the resulting motor response ramping up prior to the response being given. Strikingly, the error decoding analysis showed that the increased stimulus position coding in the latter part of the epoch reflected the participant’s decision about the stimulus more closely than the physical stimulus presented to them, providing strong evidence for the connection between specific neural responses decoded with MVPA and behaviour.

## Discussion

In this study we used MVPA with MEG to characterise the neural dynamics of stimulus, cue, rule and response coding in a difficult response-mapping task, and the link between these codes and behaviour. Our results showed a clear and orderly progression of task-relevant information coding after the stimulus was presented, while analyses aligned to the response time revealed that information coding for the stimulus and motoric response ramped up over the ∼1 second before the response was given, in a manner reminiscent of evidence accumulation (e.g., Pisauro et al., 2017; Tagliabue et al., 2019). Strikingly, for trials on which participants made an error in the stimulus position, information coding initially corresponded to physical stimulation, but later accumulated in the opposite direction, so that activity patterns at later timepoints resembled those encoding the *incorrect* stimulus. This provides a crucial demonstration that patterns of neural activity recorded and classified in this way can be predictive of behaviour. These findings give insight into the dynamics of processes underlying cognitive control and provide a clear link between neural responses and behaviour.

The difficult response-mapping task implemented in this study required complex processing for successful performance. The task involved processing different types of perceptual information (cue, stimulus), conversion of the cue into the appropriate rule, application of the relevant rule to the stimulus position, and selection of the correct button-press response. Using MEG decoding with a carefully balanced experimental design, we were able to investigate the coding of each of these types of relevant information over time, and observe the succession of the different task-related features. We summarise and consider the findings below.

### Information coding after stimulus onset: correct trials

Our results demonstrated different dynamics for perceptual, rule-related and motor processes for analyses aligned to stimulus onset (Figure 3A). Stimulus position was represented early in the time course (<100ms after stimulus onset), consistent with early retinotopic visual processes (Di Russo et al., 2005; Im et al., 2007). The cue was represented shortly thereafter at a time that is consistent with general colour processing and in line with previous work that found colour decoding was most evident from 135-155ms after image onset (Teichmann et al., 2019, 2020).

Rule information, by contrast, was most evident at around 600ms and maintained for longer than cue information, perhaps reflecting the ongoing process of combining the stimulus and rule information to derive the response. Motor responses emerged last and exhibited a broad, shallow peak, perhaps reflective of the wide range of response times in the task. Our data thus emphasise the progression of coding for different types of task-relevant information over time, despite the relevant sensory information (stimulus and cue) being presented simultaneously.

These onset-aligned analyses are consistent with previous time-resolved multivariate analyses that showed progression of task-related information after stimulus presentation (Hebart et al., 2018; Hubbard et al., 2019; Kikumoto & Mayr, 2020; Wen et al., 2019). For example, Hebart et al (2018) showed that task-related information was evident in the MEG signal shortly after a task cue was presented, but ramped up again after the target object was presented. In a more complex design, Hubbard et al (2019) used cued task-switching with electroencephalography (EEG) which allowed them to look at coding of multiple task-related aspects over time using oscillatory power in the neural signal. Like our cue and rule results, they showed cue information preceded task rule information, and task decoding was prolonged throughout the trial period. Relevant and irrelevant stimulus information was evident after the stimuli were presented, and response information was present later in the signal. Here, we show that a similar cascade of information arises when the cue and target stimulus are presented simultaneously. Our results and this previous work show that different task features are represented in the brain with different temporal dynamics, but there are time periods when multiple types of features are represented, potentially giving the neural correlates for the information integration needed on these tasks.

The cue colour decoding we observed is indicative of transient cue processing before the relevant rule was selected, at which point the colour distinctions (e.g., blue vs yellow, when both indicate rule 1) becomes irrelevant. In contrast, the prolonged coding of stimulus position and rule (far exceeding the stimulus presentation time of 500ms) likely reflects position and rule information being maintained, accumulated and/or manipulated as it is combined to achieve the correct response. Previous fMRI research has shown that the MD regions in frontoparietal cortex represent a range of information including details of stimuli and task rules (Woolgar et al., 2016), with particular emphasis on information that is task relevant (Woolgar et al 2015a, Jackson et al 2016), perceptually confusable (Woolgar et al., 2011) or difficult, like our task rules (Woolgar, Afshar, et al., 2015). The representations of stimulus position and rules we observed at late stages of processing (> 500ms after stimulus onset) would be theoretically consistent with processing within higher-level frontoparietal regions. For example, previous combined MEG-fMRI research has shown task-related enhancement of relevant features occurred after 500ms following stimulus presentation, and task coding in posterior parietal cortex and lateral prefrontal cortex seemed to peak from 500ms (Hebart et al., 2018). Moreover, attention enhances task-relevant information in frontoparietal regions from 500ms (Moerel et al., 2021). On the other hand, task-relevant stimulus information also seems to persist in occipital regions until these late timepoints (Goddard et al., 2016; Hebart et al., 2018; Moerel et al., 2021). Our exploratory sensor searchlight analyses suggested that early position, cue and rule information was evident in occipital sensors, whereas later position information was more diffuse across the brain, and later rule information was more frontal. Future work could address the spatial nature of these processes with more precision, perhaps using computational methods to combine fMRI and MEG data such as similarity-based fusion (Cichy et al., 2014).

### Information coding prior to the response: correct trials

The analyses aligned to the response time provided rich additional information about the temporal dynamics of stimulus, cue, rule and response coding (Figure 3B). We expected that these response-aligned analyses would emphasise higher-level decision-related processes required for behaviour which might not be so salient in data aligned to stimulus onset because of variability in their timing (Vidaurre et al., 2018). Previous EEG work has shown neural signals ramp up during perceptual decision-making, which has been described as evidence accumulation (e.g., Pisauro et al., 2017; Tagliabue et al., 2019), but these effects could be related to a general decision-making process rather than involving information about the stimulus of interest, and could be confounded with preparatory motor activity. Here, using decoding, we were able to assess the dynamics of different types of task-related information, separate from and in addition to response information, that was represented in the brain before the response. The results revealed an increase in stimulus information from approximately 1000ms prior to the response that peaked around the response time, a pattern which was noticeably absent in the onset-locked analyses. Response coding, by contrast, showed a later, sharper ramping in information that peaked just after the response was made. The ramping of stimulus position and response coding was, for the most part, whe n the stimulus and cue were no longer visible; the stimulus and cue were only presented for 500ms, and the median response time was over 1400ms, so on a typical trial there was no stimulus presented in the 900ms prior to the response. Therefore, instead of perceptual accumulation, these pre-response representations appear to be internally generated codes that reflect the system moving towards different end states as the person arrives at their decision.

The results revealed concurrent coding of position, rule and response information prior to the response, which might reflect the need to combine position and rule information to select the appropriate response. Kikumoto & Mayr (2020) recently investigated the temporal dynamics of action selection using EEG in a cued rule selection task and provided evidence that conjunctions between task-relevant features are necessary for action selection. In addition to the succession of individual task features, they found rule-stimulus-response conjunctive representations could be decoded using stimulus-aligned EEG, and the strength of the conjunctive information was associated with faster responses, providing a link with behaviour. Other work has used temporal decomposition of EEG data and concluded that stimulus-response bindings have different temporal profiles to stimulus information, with gradual activation and decay over time (Takacs et al., 2020). Our exploratory searchlight decoding results showed that position and rule information prior to the response was mostly lateral and diffuse across the sensors but with similar spatial patterns (though lower for rule decoding), potentially reflecting the integration of task-relevant information within brain regions. We did not explicitly set out to look at conjunctive representations, but our results certainly fit with this account. During goal-directed behaviour, it seems that multiple task-relevant features are represented concurrently, presumably reflecting the need for this information to be maintained, and are then combined over time.

Our onset-aligned and response-aligned analyses revealed complementary aspects of the data. The pattern of results suggests that onset-aligned analyses may be most sensitive to perceptual responses, while response-aligned analyses may capture processes that are time-varied relative to stimulus onset and more closely yoked to the time of response, such as higher decision and motor preparation processes. Specifically, we found that stimulus position and cue colour had sharp initial decoding when aligned to stimulus onset, which was not visible after realignment to response. However, neural representations of stimulus and response exhibited a ramping accumulation before the button was pressed, that was not visible in onset-aligned data. This highlights the utility of including both approaches, perhaps particularly for difficult tasks with substantial response time variability, to yield additional information about the dynamics underlying successful task performance.

### Information coding leading to incorrect behaviour: error trials

To test whether the neural coding of task-relevant information detected with MVPA reflects activity necessary to successfully perform a task, we examined how these codes changed when participants made errors. We focused on decoding stimulus and rule information during stimulus errors and rule errors, situations in which the decision made could be dissociated from the stimulus and rule cue presented. Stimulus errors consisted of trials on which participants correctly applied the rule but confused the stimulus. Despite the behavioural evidence for correct rule use on these trials, there was only some evidence of rule coding, perhaps reflecting weak rule coding in general (on correct trials) and the limited number of error trials. However, stimulus position coding on stimulus error trials revealed a striking result: initial stimulus coding showed some fleeting evidence of the correct stimulus neural pattern, but prior to the response, stimulus coding became consistent with the incorrect stimulus. Thus, onset-aligned analyses and responses at early timepoints reflected perception, while response-aligned analyses and coding at later timepoints reflected behaviour. Recall in the paradigm that stimulus position was designed to be confusable, and a stimulus position error, by definition, means participants confused two (out of four) stimulus positions. When the stimulus was presented, participants would see it, which is consistent with brief veridical stimulus position decoding, but the insufficient maintenance of this information correlates with the behavioural performance: participants could not localise the stimulus precisely, which led to a decision in favour of the wrong stimulus. It is this internal decision-related process that seems to be reflected in below-chance (incorrect stimulus) decoding before the response. Previous work has shown that higher, but not early, perceptual regions, reflect behaviour in terms of accuracy (Walther et al., 2009; Williams et al., 2007) and reaction time (Grootswagers et al., 2018), although none of these studies revealed the code reversal needed for a strong link with behaviour. Here, we used the temporal domain to show *what* was coded on error trials at different stages of processing. There was a dissociation between the coding of early perceptual information and the stimulus decision used to generate the behavioural response.

Rule errors consisted of trials on which participants appeared (in their behaviour) to apply the wrong rule to the correct stimulus. Accordingly, for stimulus position coding, the classifier trained on correct trials could successfully classify the stimulus position after onset and prior to the response on rule error trials. This indicates that the stimulus coding reversal seen above was diagnostic of the particular type of behavioural error, rather than reflective of errors in general, indicating a tight link with the specific decision made and reflected in behaviour.

For rule coding, we again found little evidence for rule coding on rule error trials. A couple of timepoints just prior to the response showed patterns of activity consistent with the incorrect rule, as we had predicted for a full double dissociation, but the effect was so transient that it is difficult to interpret with confidence. This may reflect the very small number of rule error trials, and/or the relatively weak coding of rule information in general in our data (potentially attributable to more variability in timing of this task aspect, and/or relatively poor signal from frontal regions that are further from the sensors in our supine MEG system). This limitation means that, in contrast to the stimulus information, we cannot conclude with confidence whether or not the rule patterns we decoded were closely linked to behaviour.

Our research contributes to the growing literature drawing links between neural responses and behaviour using MVPA. Using spatial and temporal neuroimaging, classifier prediction errors and distances from the classifier boundary have been shown to correlate with behavioural error patterns and reaction times (e.g., Carlson et al., 2014; González-García et al., 2021; Grootswagers et al., 2018; Walther et al., 2009). Here, we argue that a tighter link between brain and behaviour can be found by testing *what* is represented on error trials when an incorrect decision is made.

The results parallel fMRI work showing frontoparietal MD regions represent the correct stimulus but wrong rule during a rule error, and the correct rule but wrong stimulus during other errors (Woolgar et al., 2019). The current study extends this work by elucidating the dynamics with which the incorrect representations evolve over time, with early representations reflecting the stimuli presented, but a late gradual accumulation towards the opposite stimulus at timepoints just prior to behavioural response. We also show here that there is a dissociation in the perceptual coding (indexed by onset-aligned analyses) and high-level decision coding (indexed by response-aligned analyses) for error trials. Specifically, on stimulus errors, after a transient representation of the veridical stimulus, activity accumulated towards a pattern state reflecting the opposite and incorrect stimulus, apparently reflecting the internal generation of accumulation towards the wrong decision. This pattern was specifically diagnostic of behavioural errors attributable to stimulus misperception, as position information was coded correctly on other types of behavioural errors.

The results of this study provide new insights into how task-relevant information is processed in the human brain to allow successful goal-directed behaviour. There was a clear progression of the onset of task-relevant information in the brain, from stimulus position and cue, to rule and then response information. Complimentary response-aligned analyses, which highlight later high-level processes aligned in time to behaviour, additionally revealed dynamics of information coding resembling an accumulation of multiple types of task-relevant information. Moreover, when participants made behavioural errors, the direction of accumulation was reversed. Under these conditions, the trajectory of representation moved in the opposite direction such that the neural pattern increasingly represented the incorrect stimulus (which had not been shown) in a manner diagnostic of the subsequent behavioural choice. The data highlight the orderly but overlapping dynamics with which several task elements can be represented in brain activity. Our findings emphasise a particular role for the trajectory of information coding at later time points in determining behavioural success or failure, and demonstrate the utility of aligning neural data differently to examine high-level complex cognitive processes.

## Acknowledgements

We thank Christopher Whyte for assistance in data collection and Dr Tijl Grootswagers for helpful discussions. This work was funded by Australian Research Council Discovery Project 170101840, Australian Research Council Future Fellowship FT170100105, Medical Research Council (UK) intramural funding SUAG/052/G101400. The authors acknowledge the Sydney Informatics Hub and the University of Sydney’s high performance computing cluster Artemis for providing high performance computing resources that contributed to these research results.

